# Tumor-hepatocyte crosstalk drives a hepatic lactate-TGF-β axis of CD8^+^ T cell exhaustion and immunotherapy resistance in small-cell lung cancer liver metastases

**DOI:** 10.64898/2026.07.30.741857

**Authors:** Amira Kazi, Yingying Cao, Chirayu Mohindroo, Abhinav Joshi, Yang Zhang, Yue Huang, Chiori Tabe, Brett Schroeder, Thorkell Andersson, Ajit Kumar Sharma, Anish Thomas

## Abstract

**Purpose:** Liver metastases confer poor outcomes and attenuate the benefit of immunotherapy across solid tumors. This study investigated how the hepatic metastatic niche promotes CD8⁺ T cell dysfunction and immunotherapy resistance in small-cell lung cancer (SCLC).

**Experimental Design:** Clinical outcomes and tumor gene expression were integrated with multi-region single-cell RNA sequencing of T cells from rapid-autopsy SCLC metastases, together with spatial transcriptomics. SCLC-hepatocyte conditioned-media models were combined with stable-isotope tracing, mass spectrometry, functional and metabolic assays, and ChIP-qPCR to define mechanisms of CD8⁺ T cell suppression.

**Results:** Liver metastases were associated with inferior survival and reduced benefit from immune checkpoint blockade. Multi-region single-cell analysis showed that CD8⁺ T cells from liver metastases exhibited an exhaustion-associated state enriched for hypoxia, lactate, and TGF-β programs. SCLC-hepatocyte crosstalk generated a lactate- and TGF-β-rich microenvironment that reduced CD8⁺ T cell effector function, proximal T cell receptor signaling, glycolytic fitness, viability, and proliferation. Stable-isotope tracing demonstrated transfer and accumulation of co-culture-derived lactate in recipient CD8⁺ T cells, with limited entry into downstream pyruvate-linked pathways. Lactate accumulation was accompanied by increased H3K18 lactylation at the *PDCD1*, *LAG3*, and *TGFB1* regulatory loci. In parallel, SCLC-hepatocyte crosstalk increased paracrine TGF-β and activated canonical SMAD2 signaling in CD8⁺ T cells. TGF-β receptor inhibition restored CD8⁺ T cell proliferation. In the phase III IMpower133 cohort, a combined lactate-TGF-β transcriptional program was associated with inferior survival, most strongly in patients with liver metastases.

**Conclusions:** Tumor-hepatocyte crosstalk generates convergent lactate and TGF-β signals that drive CD8⁺ T cell dysfunction in liver metastases. This hepatic immune-metabolic circuit provides a potential mechanism for immunotherapy resistance and supports therapeutic strategies targeting TGF-β signaling in liver-metastatic SCLC.

**Translational Relevance:** Patients with SCLC liver metastases have poor outcomes and derive limited benefit from immune checkpoint blockade, but actionable mechanisms of hepatic immune resistance remain undefined. We identify an immune-metabolic circuit in which SCLC–hepatocyte crosstalk generate lactate and TGF-β signals that converge on CD8⁺ T cells. Stable-isotope tracing demonstrates the transfer and accumulation of tumor–hepatocyte-derived lactate in recipient T cells, which causes H3K18 lactylation at exhaustion- and TGFB1-associated loci. In parallel, paracrine TGF-β activates canonical SMAD signaling and reinforces proliferative dysfunction. TGF-β receptor inhibition restores CD8⁺ T cell proliferation. In the phase III IMpower133 cohort, a combined lactate–TGF-β program is associated with inferior survival, particularly among patients with liver metastases. These findings provide a mechanistic and biomarker framework for testing TGF-β-directed strategies in liver-metastatic SCLC, a population with substantial unmet clinical need.

## Introduction

Liver metastases are a major barrier to effective cancer therapy and are associated with poor survival and reduced responsiveness to immune checkpoint blockade across multiple solid tumors (1–3). The liver is uniquely positioned to shape antitumor immunity because of its physiologic tolerogenic state, high metabolic activity, and constant exposure to gut-derived antigens and inflammatory cues (4,5). Metastatic tumors can exploit these features to establish an immunosuppressive niche that promotes immune escape and therapeutic resistance.

CD8⁺ T cell exhaustion is a central mechanism of immune escape in cancer (6). Exhausted T cells progressively lose proliferative capacity, cytotoxic activity, and effector cytokine production while acquiring inhibitory receptors such as PD-1, LAG3, TIM-3, and TIGIT (7). Although persistent antigen stimulation is a principal driver of exhaustion, tissue-specific signals influence whether T cells retain functional plasticity or acquire a more durable dysfunctional state. Within the hepatic metastatic niche, metabolic stress (8,9), hypoxia (8), lactate accumulation (10,11), and suppressive cytokines (12,13) may converge on transcriptional and epigenetic programs that restrict CD8⁺ T cell function. How these pathways interact to shape T cell exhaustion in SCLC liver metastases remain poorly defined.

Small-cell lung cancer (SCLC) provides a clinically important model in which to study this problem (14,15). Nearly all patients develop early systemic dissemination (16), with frequent hepatic involvement (16–18), that portends particularly poor outcomes (19,20). Although platinum-etoposide chemotherapy and immune checkpoint blockade have modestly improved outcomes for patients with extensive-stage disease, durable responses remain uncommon (21,22). Patients with liver metastases have particularly poor outcomes and appear to derive less benefit from immunotherapy than those without hepatic involvement. Understanding how the liver metastatic microenvironment suppresses antitumor immunity could therefore reveal therapeutic vulnerabilities in this high-risk population.

Here, we integrate patient-derived single-cell transcriptomics, spatial profiling, and functional immune-metabolic studies to define the immunosuppressive mechanisms within the SCLC liver metastatic niche. We find that tumor–hepatocyte crosstalk generates two convergent immunosuppressive signals. First, SCLC–hepatocyte interactions produce lactate that is taken up and accumulated by CD8⁺ T cells, causing increased H3K18 lactylation at exhaustion- and TGFB1-associated loci. Second, SCLC–hepatocyte crosstalk increases paracrine TGF-β, which activates canonical TGF-β–SMAD signaling in CD8⁺ T cells. Together, lactate and TGF-β impair CD8⁺ T cell function and proliferation. TGF-β receptor inhibition restores proliferation. The combined lactate–TGF-β program was most strongly associated with inferior survival in patients with liver metastases. These findings define a hepatic lactate–TGF-β axis that reinforces CD8⁺ T cell exhaustion in SCLC liver metastases.

## Methods

### Patient survival and clinical outcome analysis

Clinical and survival data from the IMpower133 phase III trial were analyzed to evaluate the prognostic significance of liver metastasis in SCLC. IMpower133 enrolled patients with extensive-stage SCLC and compared first-line chemoimmunotherapy with chemotherapy alone. Linked clinical data were obtained from the European Genome-Phenome Archive (accession: EGAD00001000195; study: EGAS00001001338). Patients were stratified according to the presence or absence of liver metastases at baseline.

Progression-free survival and overall survival were estimated using the Kaplan–Meier method, and differences between groups were assessed using the log-rank test. Hazard ratios and 95% confidence intervals were calculated using Cox proportional hazards regression models. Univariable Cox regression analyses were performed to evaluate the association between individual clinical variables and survival outcomes. Variables that were clinically relevant or statistically significant in univariable analysis were subsequently included in multivariable Cox regression models to identify independent prognostic factors.

### Single-cell RNA-seq analysis

Single-cell RNA-seq data from multi-site SCLC metastases were analyzed using Seurat v4. Annotated T and natural killer cell populations were subsetted for downstream analysis. CD8⁺ T cells were defined as CD3D⁺CD8A⁺CD8B⁺CD4⁻ cells, with natural killer cells excluded based on KLRD1 and FCGR3A expression. Cells were grouped by anatomical origin into liver and non-liver groups. Data were normalized, scaled, and subjected to principal component analysis. The top 20 principal components were used for UMAP visualization and graph-based clustering. Principal component loadings were examined to assign biological programs. Differential expression between liver-derived and non-liver-derived CD8⁺ T cells was performed using the Wilcoxon rank-sum test with Seurat FindMarkers. Module-score and pseudotime distributions were compared using Wilcoxon tests. Pathway activity was quantified using Seurat AddModuleScore. Gene sets included exhaustion-associated genes, including PDCD1, HAVCR2, LAG3, and TOX; hypoxia-associated genes, including HIF1A, LDHA, and CA9; lactate-metabolism-associated genes, including LDHA, SLC16A3, and SLC2A1; and TGF-β signaling genes, including TGFBR1, TGFBR2, SMAD2, SMAD3, and SMAD4. Trajectory inference was performed using Monocle3. CD8⁺ T cells were converted to a cell_data_set object, and trajectories were learned using standard workflows. Genes were ordered by peak expression, and module-level trends were computed as mean z-scores across genes. For temporal analysis, cells were stratified into early and late pseudotime groups based on the median pseudotime value. UMAP plots, feature plots, violin plots, and pseudotime trend analyses were generated using Seurat and ggplot2. Feature plots were scaled using quantile cutoffs, and violin plots included embedded boxplots and statistical annotations.

### NicheNet ligand-activity analysis and pathway-score correlations

Ligand-receptor signaling associated with liver-infiltrating exhausted CD8⁺ T cells was inferred using the NicheNet framework. Cells were assigned to liver or non-liver groups, and differential expression in receiver T cells was computed using Seurat FindMarkers with the Wilcoxon test, a log-fold-change threshold of 0.25, and a minimum fraction of 0.1. NicheNet resources, including the ligand-target matrix, ligand-receptor network, and weighted networks, were loaded from human reference files and used to calculate ligand-activity scores for candidate upstream ligands. TGF-β-family ligands were highlighted among the predicted ligand programs. To quantify lactate-associated signaling programs, pathway and module scores were computed using curated gene sets. For sample-level analyses, scores were averaged across cells within each sample to generate pseudobulk values. Associations between lactate scores and T cell functional states, including checkpoint, effector, TGF-β, and dysfunction scores, were assessed by Spearman correlation and visualized with group-specific linear fits. An extreme liver lactate outlier identified by an interquartile-range criterion was excluded from correlation fitting but retained for display.

### CD8⁺ T cell isolation and activation

Peripheral blood apheresis products from healthy donors were obtained from the NIH Blood Bank. Peripheral blood mononuclear cells were isolated by density-gradient centrifugation using Ficoll-Paque. CD8⁺ T cells were purified from peripheral blood mononuclear cells using magnetic negative selection with the Miltenyi Biotec CD8⁺ T Cell Isolation Kit according to the manufacturer’s instructions. Purified CD8⁺ T cells were activated using Dynabeads Human T-Activator CD3/CD28 at the recommended bead-to-cell ratio. Cells were cultured in DMEM GlutaMAX supplemented with 10% fetal bovine serum in 24-well plates at a density of 1 × 10⁶ cells per well and maintained at 37°C with 5% CO₂. After 24 hours of activation, magnetic beads were removed, and cells were transferred to the indicated experimental media.

### Cell lines, culture conditions and conditioned-media preparation

The SCLC cell line DMS273 was obtained from MilliporeSigma (Cat. No. 95062830-1VL). Human immortalized hepatocytes (HHL5) were generously provided by the Arvind H. Patel laboratory (23). DMS273 and HHL5 cells were maintained in Dulbecco’s Modified Eagle Medium supplemented with 10% fetal bovine serum and 1% penicillin-streptomycin at 37°C in a humidified incubator with 5% CO₂. Cell lines were routinely tested for mycoplasma contamination using the MycoAlert Mycoplasma Detection Kit (Lonza).

For conditioned-media experiments, DMS273 cells, HHL5 cells, or DMS273/HHL5 co-cultures were grown under standard culture conditions for 4 days. Media were replenished on day 2. On day 4, conditioned media were collected, passed through 0.20-µm filters to remove cellular debris, and diluted to 40% v/v with fresh baseline medium before use in T cell assays. Baseline medium consisted of DMEM GlutaMAX supplemented with 10% fetal bovine serum. Where indicated, 10 mM sodium lactate was included as a lactate-supplemented positive-control condition.

### CD8⁺ T conditioned-media exposure

After 24 hours of activation, magnetic beads were removed, and cells were transferred to the indicated experimental media. CD8⁺ T cells were maintained in the indicated media conditions for 5 days before downstream analyses, including flow cytometry, immunoblotting, extracellular flux analysis, lactate quantification, viability, and proliferation assays. This duration was chosen to allow sustained exposure to soluble factors present in the conditioned media and to capture coordinated changes in cytokine production, inhibitory receptor expression, TCR signaling, metabolic activity, and proliferation, consistent with prior studies showing that T cell dysfunction develops through persistent stimulation and tumor microenvironmental stress (7,8,24).

### Flow cytometry

After 5 days of culture in conditioned media, CD8⁺ T cells were stimulated to assess cytokine production. Cells were incubated with phorbol 12-myristate 13-acetate at 50 ng/mL, ionomycin at 1 µg/mL, and brefeldin A at 5 µg/mL for 4–6 hours at 37°C to induce cytokine expression and block secretion. Cells were collected, washed three times with flow cytometry buffer consisting of PBS with 2% fetal bovine serum, and incubated with human serum to block Fc receptors. Surface staining was performed with antibodies against CD69 and CD25 as activation markers and PD-1 and LAG3 as dysfunction-associated markers. For intracellular cytokine detection, cells were fixed and permeabilized using permeabilization buffer according to the manufacturer’s instructions and stained for IFN-γ and TNF-α. Following staining, cells were washed and resuspended in PBS for analysis on a flow cytometer. Data were acquired using standard acquisition settings and analyzed using FlowJo software.

### Immunoblotting

For immunoblotting, CD8⁺ T cells were collected after 5 days of culture under the indicated media conditions. Approximately 5 × 10⁶ cells per sample were washed with cold PBS and lysed in 250 µL of 2× Laemmli sample buffer (Bio-Rad, Cat. No. 1610737) supplemented with 5% 2-mercaptoethanol (Sigma, Cat. No. M3148-100ML). Lysates were sonicated using five cycles of 30 seconds on/off at 4°C, vortexed, boiled for 5 minutes, and centrifuged at 11,000 × g for 5 minutes at 4°C. Supernatants were collected, and equal amount of proteins from each sample was resolved by SDS-PAGE using 10% gels for total protein analyses and 15% gels for histone analyses. Proteins were then transferred to PVDF membranes (Millipore, Cat. No. IPVH304F0).

Membranes were blocked in 5% non-fat dry milk in 0.1% TBS-T, consisting of 10 mM Tris-HCl pH 7.6, 150 mM NaCl, and 0.1% Tween-20, for 1 hour at room temperature. Membranes were incubated overnight at 4°C with primary antibodies diluted in 1% BSA in 0.1% TBS-T. After washing, membranes were incubated with appropriate HRP-conjugated mouse or rabbit secondary antibodies diluted in 5% milk for 1 hour at room temperature. Blots were developed using Immobilon Western or SuperSignal West Femto chemiluminescent substrates and imaged using either film-based darkroom exposure or a chemiluminescence imaging system.

### Seahorse metabolic analysis

Extracellular acidification rate was measured using a Seahorse XFe/XF96 Analyzer. Seahorse XF96 microplates were coated with poly-L-lysine for 1 hour at room temperature to facilitate attachment of CD8⁺ T cells. After removal of the coating solution, plates were dried for 1-3 hours at 37°C. Seahorse XF cartridges were hydrated overnight in Seahorse XF calibrant solution at 37°C in a non-CO₂ incubator. After 5 days of culture under the indicated media conditions, activated CD8⁺ T cells were counted and seeded into poly-L-lysine-coated wells at 2 × 10⁵ cells per well. Cells were equilibrated for 60 minutes at 37°C in a non-CO₂ incubator in Seahorse XF assay medium consisting of Seahorse XF Base Medium supplemented with 10 mM glucose, 2 mM glutamine, and 1 mM pyruvate.

Basal ECAR was measured followed by a sequential injection of compounds including oligomycin, FCCP, and rotenone/antimycin A, according to the manufacturer’s instructions. ECAR were used to assess overall bioenergetic fitness of CD8⁺ T cells following exposure to the indicated conditioned-media conditions. Data were normalized to the number of cells seeded per well.

### Lactate quantification

Lactate levels in conditioned media and intracellular fractions were quantified using the Elabscience Bioluminescent Lactate Assay Kit according to the manufacturer’s instructions. For conditioned media, samples were collected on day 4 of co-culture and clarified by centrifugation at 500 × g for 5 minutes to remove cells and debris. Samples were analyzed directly or diluted as needed to fall within the assay’s dynamic range.

For intracellular lactate measurements, CD8⁺ T cells were harvested, washed twice with cold PBS, and lysed using the cell lysis buffer provided with the assay kit. Lysates were clarified by centrifugation at 12,000 × g for 5 minutes at 4°C, and supernatants were collected for analysis. Protein concentration was determined using a BCA assay to normalize intracellular lactate levels.

For both media and intracellular samples, 20 µL of sample or standard was added to a 96-well plate in technical triplicate. Reaction mix containing the bioluminescent lactate detection reagents was added, and plates were incubated at room temperature for 10–20 minutes in the dark. Luminescence was measured using a plate reader. Lactate concentrations were calculated from standard curves and reported as micromolar concentrations for media or normalized to total protein for intracellular samples.

### Stable-isotope tracing and metabolite-labeling analysis

DMS273 cells and HHL5 hepatocytes were seeded together in T75 flasks at a 4:1 ratio and maintained in DMEM GlutaMAX supplemented with 10% FBS. After 48 hours, the medium was replaced with glucose-tracer medium containing uniformly labeled ^13^C-glucose ([U-^13^C_6_] glucose), and the co-cultures were incubated for an additional 48 hours. This yielded a total co-culture duration of 4 days. Conditioned medium was then collected, filtered to remove cells and debris, and diluted to a final concentration of 40% in T cell culture medium.

CD8⁺ T cells isolated from healthy-donor PBMCs were activated in parallel for 24 hours in 24-well plates at 1×10^6^ cells per well using 10 µL of CD3/CD28 activation beads per well. The activation beads were subsequently removed, and the T cells were cultured for 5 days in the indicated media at 5×10^6^ cells per well. Three conditions were analyzed: inactive CD8⁺ T cells cultured in medium containing unlabeled glucose, activated CD8⁺ T cells cultured in 40% conditioned medium from DMS273/HHL5 co-cultures maintained with unlabeled glucose, and activated CD8⁺ T cells cultured in 40% conditioned medium from DMS273/HHL5 co-cultures maintained with [U-^13^C_6_] glucose. At the end of the incubation period, T cells were collected by centrifugation, and the resulting cell pellets were stored at −80°C until metabolite-labeling analysis.

Metabolic flux analysis using was performed by Human Metabolome Technologies, Inc. Three biological replicates for each condition were used for this experiment. The samples were analyzed using capillary electrophoresis time-of-flight mass spectrometry (CE-TOFMS, Agilent Technologies) in two modes to detect both anionic and cationic metabolites. Detected peaks were then extracted using MasterHands ver. 2.17.1.11 to obtain m/z, migration time (MT), and peak area. Putative metabolites were assigned based on HMT’s target library and their isotopic ions using m/z and MT. Absolute quantitation was performed for the total amount of each detected metabolite.

Isotopologue enrichment was assessed in lactate and metabolites associated with pyruvate-linked metabolism. Lactate m+3 was measured as the primary indicator of labeled lactate acquisition, and alanine m+3 was evaluated as an indicator of labeling within the pyruvate-associated metabolite pool. m+2 isotopologues of 2-oxoglutarate, glutamate, malate, and aspartate were examined as downstream labeling associated with pyruvate dehydrogenase–mediated carbon entry. Malate m+3 and aspartate m+3, together with downstream 2-oxoglutarate m+3 and glutamate m+3, were examined as labeling associated with pyruvate carboxylase–mediated entry. Potential carryover and utilization of residual [U-^13^C_6_] glucose were assessed by measuring glucose 1-phosphate m+6, glucose 6-phosphate m+6, fructose 6-phosphate m+6, fructose 1,6-bisphosphate M+6, and glyceraldehyde 3-phosphate M+3.

### CD8⁺ T cell viability assay

CD8⁺ T cell viability under the indicated culture conditions was assessed using the ATPLite Luminescence Assay Kit according to the manufacturer’s protocol. Cells were seeded into 96-well plates at a density of 1 × 10⁵ cells per well in normal media or conditioned media. Cells were incubated under standard culture conditions at 37°C with 5% CO₂ for 5 days. At the end of the incubation, ATPLite reagent was added to each well, and luminescence was measured using a plate reader according to the kit instructions. Luminescence values were normalized to untreated controls to determine relative cell viability. Assays were performed in technical triplicate for each condition.

### CD8⁺ T cell proliferation assay

CD8⁺ T cell proliferation under the indicated culture conditions was monitored using the IncuCyte live-cell imaging system. Cells were seeded into 96-well plates at a density of 1 × 10⁵ cells per well in normal media or conditioned media. Cells were cultured under standard conditions at 37°C with 5% CO₂ for 5 days. Phase-contrast images were acquired automatically every 6 hours using the IncuCyte system analyzed using IncuCyte software.

### Quantification of TGF-β1 and TGF-β2 by ELISA

TGF-β1 and TGF-β2 levels in conditioned media were quantified using commercially available ELISA kits according to the manufacturers’ instructions. TGF-β1 was measured using the Quantikine ELISA kit from R&D Systems, and TGF-β2 was measured using a human/mouse/rat TGF-β2 ELISA kit from Antibodies.com. Conditioned media were collected from DMS273, HHL5, and DMS273/HHL5 co-culture systems after 4 days of incubation. Samples were centrifuged at 500 × g for 5 minutes to remove cellular debris and stored at −80°C until analysis. Before ELISA, samples were thawed on ice and diluted as needed to fall within the linear range of the assay. To measure total TGF-β levels, samples were acid-activated according to the manufacturer’s protocol to convert latent TGF-β to its active form. Standards and samples were added to 96-well plates in technical triplicate, followed by incubation with capture and detection antibodies. After washing, substrate solution was added, and absorbance was measured at 450 nm using a microplate reader. Concentrations of TGF-β1 and TGF-β2 were calculated from standard curves.

### ChIP-qPCR

Cells were crosslinked in 1% formaldehyde for 10 minutes at room temperature with gentle agitation to preserve protein-DNA interactions. Crosslinking was quenched with 125 mM glycine for 5 minutes. Cells were washed twice with cold PBS and pelleted. Cell pellets were lysed using ChIP lysis buffer containing protease inhibitors. Nuclei were isolated, and chromatin was sheared by sonication to obtain fragments of approximately 200–500 bp. Equal amounts of chromatin were incubated overnight at 4°C with anti-H3K18 lactylation antibody or normal IgG control.

Antibody-chromatin complexes were captured using Protein A magnetic beads for 2–4 hours at 4°C. Beads were sequentially washed with low-salt, high-salt, LiCl, and TE buffers. Chromatin was eluted from beads, and crosslinks were reversed by incubation at 65°C overnight. DNA was purified using a Qiagen Spin Miniprep Kit and quantified using NanoDrop. Enrichment of H3K18 lactylation at promoter regions was assessed by quantitative PCR using primers targeting PDCD1, LAG3, and TGFB1 promoters. Primers were designed to amplify regions near transcription start sites. qPCR reactions were performed using SYBR Green master mix on a real-time PCR system.

### Statistical analysis

Statistical analyses were performed using GraphPad Prism 10.4.1 and appropriate statistical software for survival modeling. Comparisons between two groups were conducted using two-sided Wilcoxon rank-sum tests or unpaired two-tailed Student’s t tests, as indicated. Survival outcomes were assessed using Kaplan-Meier estimates and Cox proportional hazards models. A P value < 0.05 was considered statistically significant.

## Results

### Liver metastases define a poor-prognosis SCLC subgroup with attenuated benefit from immunotherapy

To define the clinical impact of hepatic metastases in SCLC, we first analyzed patients from the phase III IMpower133 trial, which established atezolizumab plus platinum-etoposide as a first-line standard for extensive-stage SCLC **(Fig. 1A)**. Among 132 patients in the immunotherapy arm, 75 had no liver metastases and 57 had liver metastases at baseline. Patients with liver metastases had significantly shorter progression-free survival than those without hepatic involvement (median PFS, 4.31 vs. 5.56 months; HR, 1.95; 95% CI, 1.30-2.91; P=0.001; **Fig. 1B**). Overall survival was also significantly reduced in patients with liver metastases (median OS, 9.32 vs. 12.22 months; HR, 1.93; 95% CI, 1.30-2.87; P=0.0009; **Fig. 1C**).

**Fig. 1:**
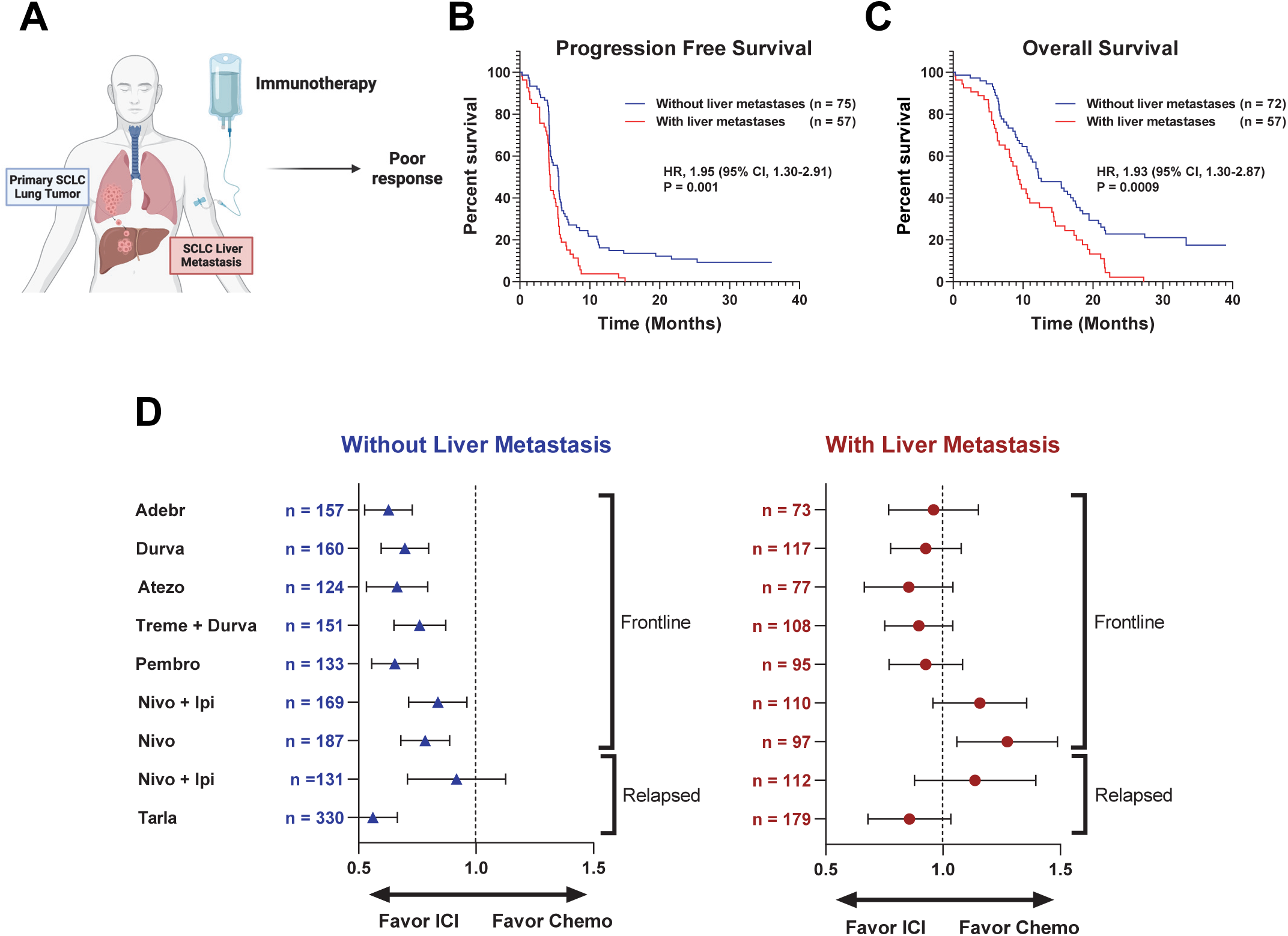
Liver metastases are associated with poor clinical outcomes and reduced benefit from immunotherapy in SCLC. **(A)** Schematic illustrating the association between liver metastases and poor response to immunotherapy in patients with SCLC. **(B and C)** Kaplan-Meier curves comparing progression-free survival **(B)** and overall survival **(C)** in patients with and without liver metastases. Hazard ratios, 95% confidence intervals, and P values are shown. **(D)** Forest plots summarizing treatment-effect estimates stratified by the presence or absence of liver metastases across frontline and relapsed SCLC clinical trials. Symbols indicate hazard ratios and horizontal lines indicate 95% confidence intervals; sample sizes are shown for each subgroup. Hazard ratios below 1 favor immune checkpoint inhibitor–based therapy, whereas values above 1 favor chemotherapy. **Abbreviations:** Adebr, adebrelimab; Atezo, atezolizumab; CI, confidence interval; Durva, durvalumab; HR, hazard ratio; ICI, immune checkpoint inhibitor; Ipi, ipilimumab; Nivo, nivolumab; Pembro, pembrolizumab; SCLC, small cell lung cancer; Tarla, tarlatamab; Treme, tremelimumab.

Because this was a post hoc analysis, the prognostic significance of liver metastasis was further evaluated after adjustment for baseline clinical and disease-related variables. In multivariable models including age, sex, smoking status, and tumor burden, liver metastasis remained associated with inferior PFS (HR, 1.68; 95% CI, 1.06–2.66) and OS (HR, 1.79; 95% CI, 1.15– 2.79), supporting hepatic involvement as an adverse prognostic factor in SCLC **(Supplementary Fig. S1A and S1B)**.

The clinical impact of liver metastasis was then examined across additional immune-based treatment strategies in SCLC. Across clinical trial datasets evaluating PD-1, PD-L1, and CTLA-4 checkpoint blockade, as well as the DLL3-targeted bispecific T cell engager tarlatamab, patients with liver metastases generally had poorer outcomes than those without hepatic involvement **(Fig. 1D)**. These data indicate that liver metastasis marks a clinically adverse SCLC subgroup with attenuated benefit from immune-based therapy.

### CD8⁺ T cells in SCLC liver metastases exhibit an exhausted, hypoxic, and lactate-associated state

To determine how the hepatic metastatic microenvironment shapes CD8⁺ T cell function in SCLC, we performed single-cell RNA sequencing of rapid-autopsy tumors collected from the liver, lung, adrenal gland, lymph node, hilum, mediastinum, pericardial mass, and skin (**Fig. 2A**). T and natural killer (NK) cells showed partial segregation by anatomical site, suggesting that the local tissue environment shapes their transcriptional states (**Fig. 2B**) (25,26). CD8⁺ T cells were identified by CD8A expression and reclustered to compare cells from liver metastases with those from non-liver sites (**Fig. 2C**). Liver-derived and non-liver-derived CD8⁺ T cells occupied distinct regions of the UMAP, indicating marked transcriptional differences between these populations (**Fig. 2D**).

**Fig. 2:**
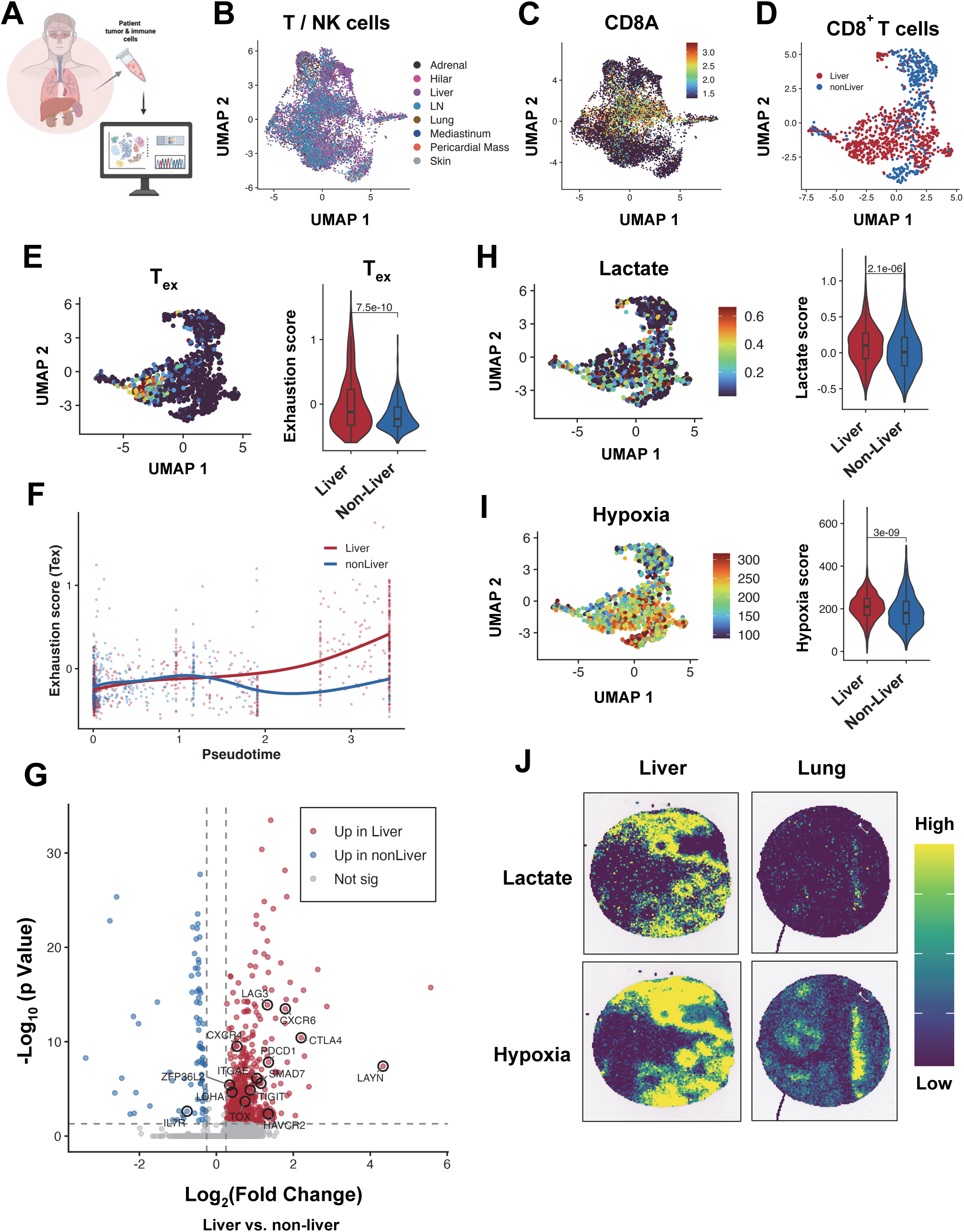
CD8⁺ T cells in SCLC liver metastases exhibit increased exhaustion, lactate, and hypoxia programs. **(A)** Schematic of multi-region single-cell RNA-sequencing analysis of rapid-autopsy SCLC tumors and associated immune-cell populations. Samples were obtained from liver, lung, adrenal gland, lymph node, hilum, mediastinum, pericardial mass, and skin metastases. **(B)** UMAP visualization of T and natural killer (NK) cells, colored by anatomical site of origin. **(C)** CD8A expression within the T- and NK-cell compartment, used to identify the CD8⁺ T cell population for downstream analysis. **(D)** UMAP visualization of CD8⁺ T cells, colored by liver versus non-liver metastatic origin. **(E)** UMAP projection of the T cell exhaustion module score and violin plots comparing exhaustion scores in CD8⁺ T cells from liver and non-liver metastases. *P* values are shown. **(F)** Exhaustion scores across inferred pseudotime in liver- and non-liver-derived CD8⁺ T cells. Points represent individual cells and curves represent smoothed trajectories. **(G)** Volcano plot of differentially expressed genes in CD8⁺ T cells from liver versus non-liver metastases. Genes enriched in liver and non-liver metastases are shown in red and blue, respectively; selected genes associated with T cell dysfunction, exhaustion, and tissue retention are labeled. Dashed lines indicate the significance and fold-change thresholds. **(H, I)** UMAP projections and violin plots showing lactate-associated (H) and hypoxia-associated (I) module scores in CD8⁺ T cells from liver versus non-liver metastases. *P* values are shown. **(J)** Spatial transcriptomic visualization of lactate- and hypoxia-associated signature scores in representative SCLC liver and lung metastases. Colors indicate relative signature intensity from low to high. **Abbreviations:** NK, natural killer; SCLC, small cell lung cancer; T_ex_, exhausted T cell; UMAP, Uniform Manifold Approximation and Projection.

Liver-derived CD8⁺ T cells had significantly higher exhaustion scores than their non-liver counterparts (**Fig. 2E**). Pseudotime analysis revealed progressive divergence of the two populations, with liver-derived cells acquiring higher exhaustion scores along the inferred trajectory (**Fig. 2F**). Differential expression analysis further showed that liver-derived CD8⁺ T cells upregulated multiple genes associated with terminal dysfunction and tissue retention, including LAG3, CTLA4, PDCD1, TIGIT, CXCR6, LAYN, TOX, and ITGAE (**Fig. 2G; Supplementary Fig. S2A).**

The liver contains physiologic porto-central gradients in oxygenation and metabolism(27,28), which hepatic tumors may intensify through local hypoxia, glycolysis, and lactate production (29,30). Because hypoxia and lactate can impair CD8⁺ T cell effector function and antitumor immunity (10,11), we asked whether liver-associated CD8⁺ T cell exhaustion coincided with transcriptional evidence of metabolic stress. Liver-derived CD8⁺ T cells had significantly higher lactate-associated and hypoxia-associated scores than cells from non-liver sites (**Fig. 2H and I**). Spatial transcriptomics confirmed the extensive lactate- and hypoxia-associated transcriptional activity in SCLC liver metastases (**Fig. 2J**). Together, these findings show that CD8⁺ T cells in liver metastases exhibit an exhaustion-associated state within a hypoxic, lactate-enriched microenvironment.

### SCLC-hepatocyte crosstalk suppresses CD8⁺ T cell effector function

The exhausted CD8⁺ T cell phenotype observed in SCLC liver metastases was associated with prominent lactate- and hypoxia-related programs. To determine whether soluble factors produced through tumor–hepatocyte interactions could recapitulate this phenotype, primary human CD8⁺ T cells were activated with CD3/CD28 beads and cultured for 120 hours in control medium or conditioned medium from DMS273 SCLC cells, HHL5 hepatocytes, or DMS273/HHL5 co-cultures (**Fig. 3A; Supplementary Fig. S3A**). Unstimulated and CD3/CD28-stimulated T cells were used as baseline and activation controls. Lactate-supplemented medium was included as a positive control because of the lactate-associated program detected in liver-derived CD8⁺ T cells (Fig. 2) and the suppressive effects of lactate on T cell function (10,11).

**Fig. 3:**
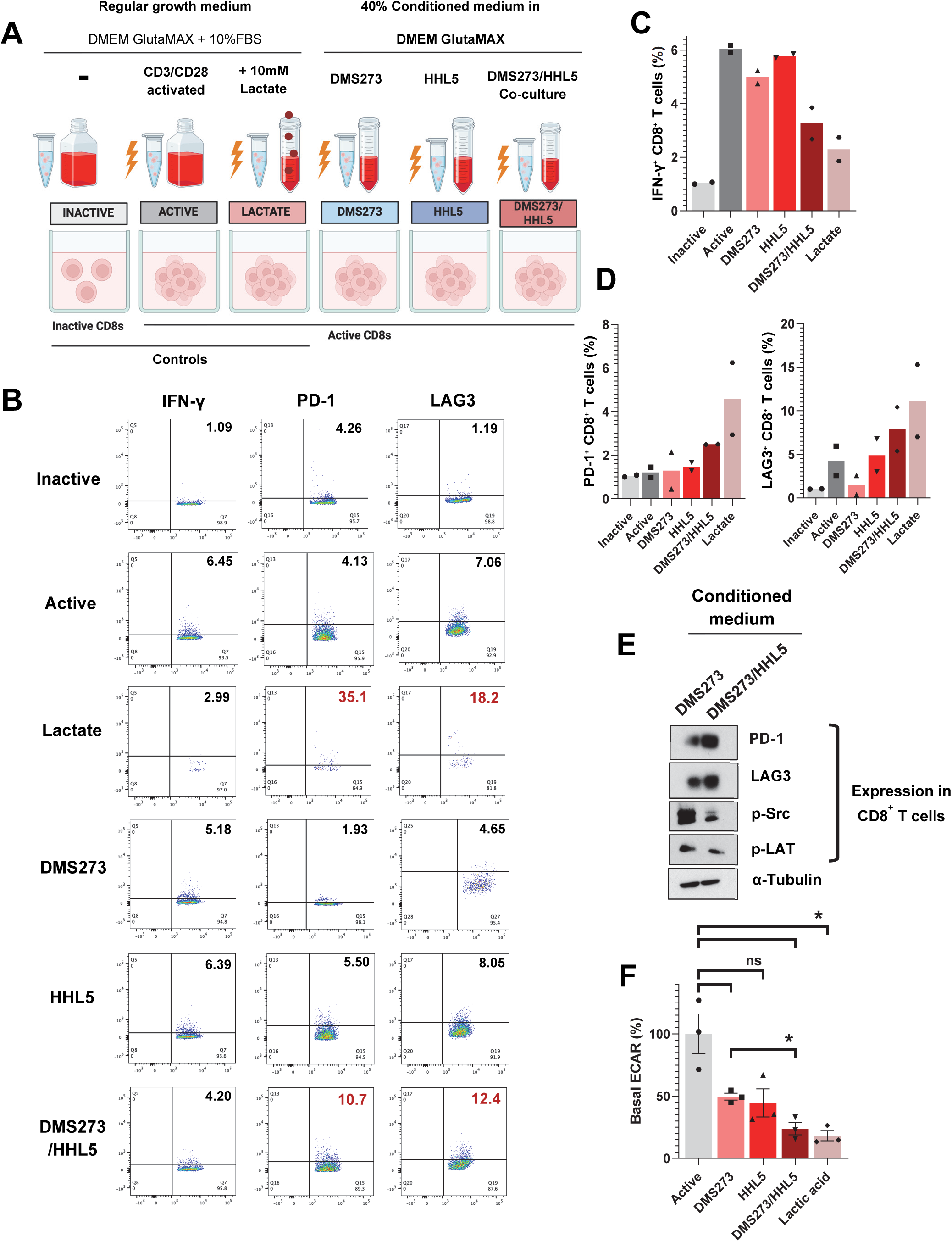
SCLC-hepatocyte-conditioned signals impair CD8⁺ T cell function. **(A)** Schematic of the experimental conditions used to evaluate CD8⁺ T cell responses. Primary human CD8⁺ T cells were activated with CD3/CD28 beads and cultured for 120 hours in control medium or medium containing 40% conditioned medium from DMS273 cells, HHL5 hepatocytes, or DMS273/HHL5 co-cultures. Unstimulated and CD3/CD28-stimulated T cells served as baseline and activation controls respectively and medium supplemented with 10 mmol/L lactic acid served as a positive control. **(B)** Representative flow-cytometry dot plots showing IFN-γ, PD-1, and LAG3 expression in CD8⁺ T cells under the indicated conditions. Values indicate the percentage of marker-positive CD8⁺ T cells. **(C and D)** Quantification of IFN-γ-positive CD8⁺ T cells as a measure of effector function **(C)** and PD-1-positive and LAG3-positive CD8⁺ T cells as exhaustion-associated populations **(D)** under the indicated conditions. Data are presented as the percentage of marker-positive CD8⁺ T cells, with inactive-control normalized across experiments to account for baseline variation. Bars represent the mean of two biological replicates (n=2), with individual biological replicates shown. **(E)** Immunoblot analysis of PD-1, LAG3, phosphorylated SRC, and phosphorylated LAT in CD8⁺ T cells cultured with conditioned medium from DMS273 cells or DMS273/HHL5 co-cultures. α-Tubulin served as the loading control. **(F)** Basal ECAR in activated CD8⁺ T cells cultured in regular activation medium, conditioned medium from DMS273 cells, HHL5 hepatocytes, or DMS273/HHL5 co-cultures, or medium supplemented with 10 mmol/L lactic acid. T cells cultured in regular activation medium exhibited the highest basal ECAR, an indicative of maximal glycolytic function, whereas lactic acid and DMS273/HHL5 co-culture-conditioned medium produced the greatest suppression (mean ± SEM, n=3). ns, not significant; ∗, *P* < 0.05. **Abbreviations:** ECAR, extracellular acidification rate; IFN-γ, interferon gamma; LAG3, lymphocyte-activation gene 3; LAT, linker for activation of T cells; PD-1, programmed cell death protein 1; SRC, SRC proto-oncogene, non-receptor tyrosine kinase.

Flow cytometry was used to measure IFN-γ as an indicator of effector activity (31) and PD-1 and LAG3 as markers of T cell dysfunction (32) (**Fig. 3B**). As expected, CD3/CD28 stimulation increased IFN-γ production. Lactate supplementation reduced IFN-γ and increased PD-1 and LAG3. A similar pattern was observed with DMS273/HHL5 co-culture-conditioned medium, which produced the largest reduction in IFN-γ and the strongest induction of PD-1 and LAG3 among the conditioned-media groups. Conditioned medium from DMS273 or HHL5 cells alone had more modest effects. These changes were reproduced across two independent experiments (**Fig. 3C and D**).

Immunoblotting further showed increased PD-1 and LAG3 expression in CD8⁺ T cells exposed to DMS273/HHL5-conditioned medium compared with DMS273-conditioned medium (**Fig. 3E**). Phosphorylation of SRC-family kinases and LAT, key proximal mediators of T cell receptor signaling, was concurrently reduced (33), indicating impaired signal propagation downstream of TCR engagement.

Because activated CD8⁺ T cells depend on glycolytic reprogramming to sustain proliferation, cytokine production, and effector function (34,35), extracellular acidification rate was measured as an index of glycolytic activity. Basal extracellular acidification was most strongly reduced in CD8⁺ T cells exposed to DMS273/HHL5-conditioned or lactate-supplemented medium relative to activated controls (**Fig. 3F; Supplementary Fig. S3B**). These findings indicate that soluble factors generated by SCLC–hepatocyte crosstalk impair CD8⁺ T cell signaling, metabolism, and effector function.

### SCLC-hepatocyte-derived lactate accumulates in CD8⁺ T cells and promotes histone lactylation

The similar effects of DMS273/HHL5-conditioned medium and lactate supplementation on CD8⁺ T cell function suggested that lactate might contribute to the suppressive activity of the co-culture medium (**Fig. 4A)**. Consistent with this, DMS273/HHL5 co-cultures produced more lactate than either cell type alone, reaching concentrations close to those in lactate-supplemented medium (**Fig. 4B**).

**Fig. 4:**
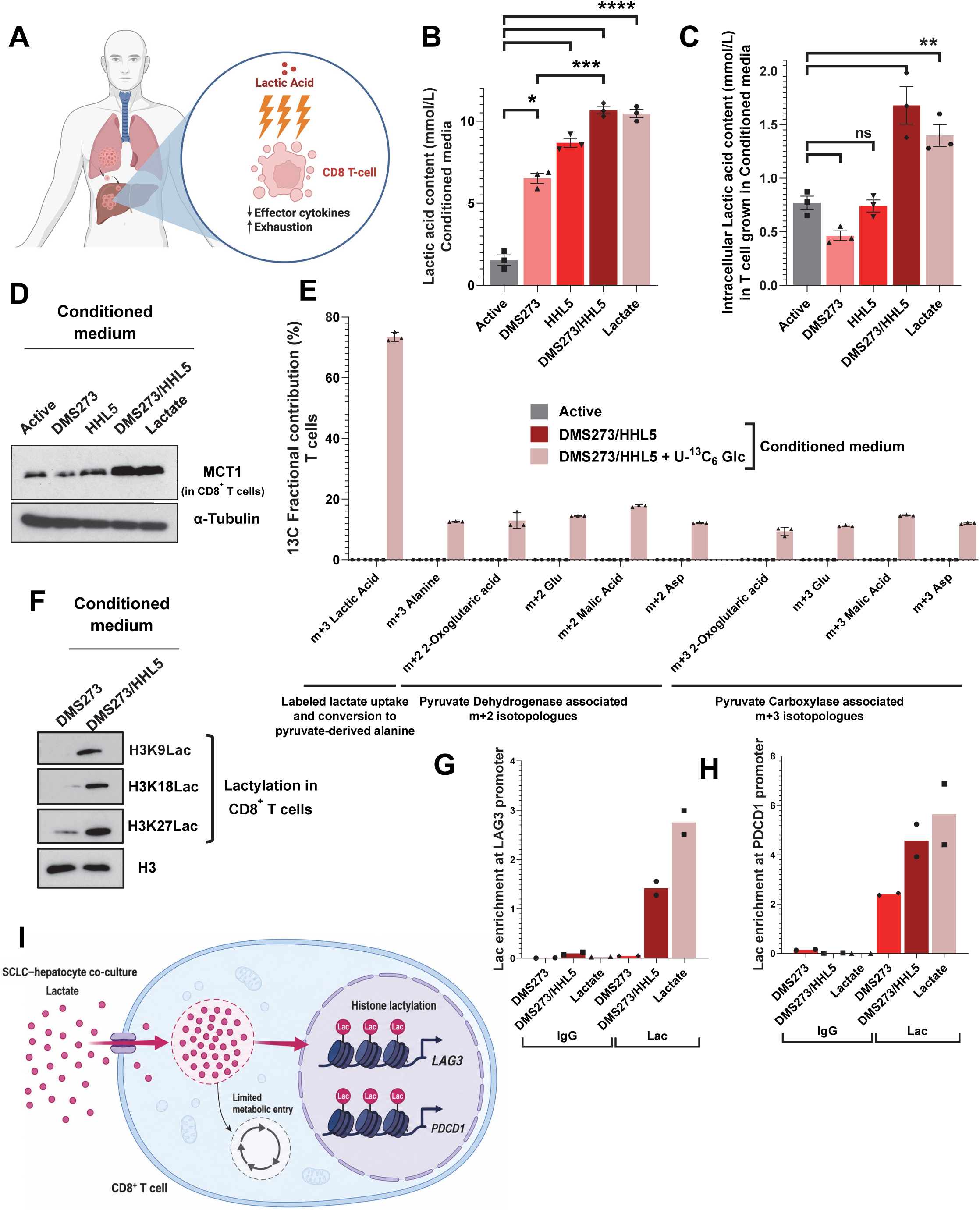
SCLC-hepatocyte-derived lactate promotes exhaustion-associated histone lactylation in CD8⁺ T cells. **(A)** Conceptual schematic illustrating the proposed contribution of lactate within the hepatic metastatic niche to reduced effector cytokine production and increased exhaustion in CD8⁺ T cells. **(B)** Lactate concentrations in regular activation medium, conditioned medium from DMS273 cells, HHL5 hepatocytes, or DMS273/HHL5 co-cultures, and medium supplemented with 10 mmol/L lactic acid. **(C)** Intracellular lactate concentrations in CD8⁺ T cells cultured under the indicated conditions. In B and C, bars represent the mean ± SEM (n = 3). Statistical significance in B and C is indicated in the figure; ns, not significant; *, P < 0.05; **, P < 0.01; ***, P < 0.001; ****, P < 0.0001. **(D)** Immunoblot analysis of MCT1 in CD8⁺ T cells cultured in regular activation medium or conditioned medium from DMS273 cells, HHL5 hepatocytes, or DMS273/HHL5 co-cultures, or with 10 mmol/L lactic acid. α-Tubulin was used as the loading control. **(E)** Fractional contribution of specific isotopologues of ^13^C-labeled metabolites in activated CD8⁺ T cells cultured in regular unlabeled activation medium, unlabeled DMS273/HHL5 co-culture-conditioned medium, or DMS273/HHL5 co-culture-conditioned medium generated with [U-13C6] glucose (mean ± SEM, n=3). **(F)** Immunoblot analysis of H3K9, H3K18, and H3K27 lactylation in CD8⁺ T cells cultured with conditioned medium from DMS273 cells or DMS273/HHL5 co-cultures. Total histone H3 served as the loading control. **(G and H)** ChIP-qPCR analysis of lactylation enrichment at the *LAG3* (G) and *PDCD1* (H) promoters in CD8⁺ T cells cultured with DMS273-conditioned medium, DMS273/HHL5 co-culture-conditioned medium, or 10 mmol/L lactic acid. Bars represent the mean of two biological replicates (n=2). **(I)** Conceptual model illustrating the uptake and intracellular accumulation of lactate produced through SCLC-hepatocyte niche, its limited entry into central carbon metabolism, and utilization of the intracellular lactate pool for histone lactylation at the *LAG3* and *PDCD1* loci in CD8⁺ T cells **Abbreviations:** ECAR, extracellular acidification rate; IFN-γ, interferon gamma; LAG3, lymphocyte-activation gene 3; LAT, linker for activation of T cells; PD-1, programmed cell death protein 1; SRC, SRC proto-oncogene, non-receptor tyrosine kinase.

Lactate enters immune cells through monocarboxylate transporters and can promote dysfunctional T cell states (36,37). CD8⁺ T cells cultured in DMS273/HHL5-conditioned medium accumulated significantly more intracellular lactate than activated controls, reaching levels comparable to those observed with lactate supplementation (**Fig. 4C**). MCT1/SLC16A1 expression increased in parallel and was highest under these same conditions (**Fig. 4D**), consistent with enhanced lactate uptake.

To determine whether CD8⁺ T cells directly acquire lactate produced by tumor-hepatocyte co-cultures, we performed an orthogonal stable-isotope tracing experiment. DMS273/HHL5 cells were cultured with U-^13^C-glucose, allowing newly produced m+3 ^13^C-lactate to be traced after transfer of the conditioned medium to CD8⁺ T cells. Two control conditions were included: inactive CD8⁺ T cells cultured with unlabeled glucose and activated CD8⁺ T cells exposed to conditioned medium from co-cultures grown with unlabeled glucose.

m+3 lactate showed the highest fractional enrichment in CD8⁺ T cells exposed to labeled conditioned medium and was undetectable in either control condition (**Fig. 4E**). By comparison, m+3 alanine was far less enriched, suggesting limited equilibration of the transferred lactate-derived carbon with the intracellular pyruvate pool. Entry into mitochondrial metabolism through pyruvate dehydrogenase (PDH)-mediated acetyl-CoA production or pyruvate carboxylase (PC)-dependent anaplerosis (38–40) was also modest. PDH-associated m+2 labeling of 2-oxoglutarate, glutamate, malate, and aspartate was detectable, whereas PC-associated m+3 labeling of malate and aspartate and downstream labeling of 2-oxoglutarate and glutamate remained low relative to m+3 lactate. These findings indicate that conditioned-medium-derived lactate accumulated predominantly within CD8⁺ T cells, with comparatively limited conversion to pyruvate and entry into the tricarboxylic acid cycle.

Because the conditioned medium was generated from co-cultures grown with U-^13^C-glucose rather than supplemented U-^13^C-lactate, residual labeled glucose could potentially have been taken up directly by recipient T cells and converted to intracellular m+3 lactate. Such a mechanism would be expected to produce prominent m+6 labeling of upper glycolytic intermediates. This labeling was minimal, arguing against glucose carryover as the principal source of intracellular m+3 lactate (**Supplementary Fig. S4A**). The labeling pattern therefore supports transfer and accumulation of co-culture-derived lactate, with comparatively limited downstream metabolic processing.

The limited downstream metabolism of imported lactate raised the possibility that its accumulation contributes to CD8⁺ T cell dysfunction through epigenetic effects. Given that lactate uptake can reinforce exhausted T cell states (36) and promote histone lactylation (41), histone lactylation was examined in CD8⁺ T cells exposed to co-culture-conditioned medium. DMS273/HHL5-conditioned medium increased H3K9, H3K18, and H3K27 lactylation relative to DMS273-conditioned medium (**Fig. 4F**). Mass spectrometry independently confirmed increased H3K18 lactylation after exposure to DMS273/HHL5-conditioned medium (**Supplementary Fig. S4B**).

H3K18la occupancy at exhaustion-associated genes was then assessed by ChIP–qPCR, with lactate-supplemented medium serving as a positive control. DMS273/HHL5-conditioned medium increased H3K18la enrichment at the LAG3 and PDCD1 promoters, closely recapitulating the pattern observed with lactate supplementation (**Fig. 4G and H**). The same conditioned medium reduced CD8⁺ T cell viability and proliferation to levels comparable to the lactate control (**Supplementary Fig. S4C and D**). These findings suggest that SCLC-hepatocyte-derived lactate accumulates in CD8⁺ T cells and is accompanied by histone lactylation at exhaustion-associated loci, providing a potential epigenetic link between the hepatic metabolic niche and T cell dysfunction (**Fig. 4I)**.

### Lactate and paracrine TGF-β converge to suppress CD8⁺ T cell proliferation

The enrichment of lactate- and hypoxia-associated programs in liver-derived CD8⁺ T cells suggested that additional signals within the hepatic niche might cooperate with metabolic stress to promote dysfunction. NicheNet ligand-activity analysis identified TGF-β-family ligands among the highest-ranked predicted regulators of the liver-enriched CD8⁺ T cell program (**Fig. 5A**). This finding was consistent with the established roles of TGF-β in hepatic immune tolerance, liver metastasis, and suppression of antitumor CD8⁺ T cell function (5,12,13,42). Gene set enrichment analysis further showed increased TGF-β pathway activity in SCLC liver metastases (**Fig. 5B**), and liver-derived CD8⁺ T cells had higher TGF-β pathway scores than cells from non-liver sites (**Fig. 5C**).

**Fig. 5:**
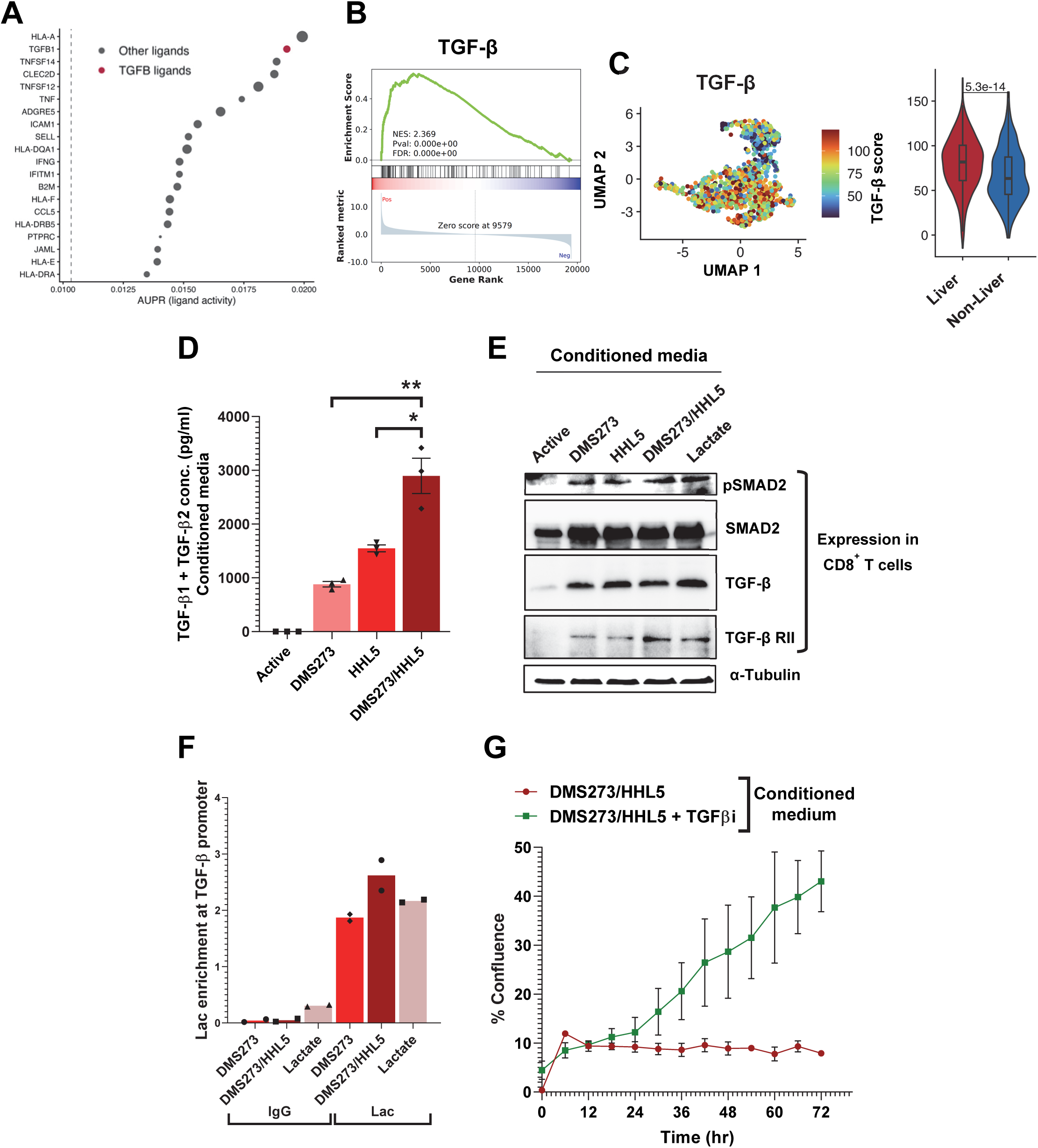
A lactate-TGF-β axis suppresses CD8⁺ T cell proliferation. **(A)** NicheNet ligand-activity analysis identifying candidate signals associated with the transcriptional program of CD8⁺ T cells from liver metastases. TGF-β family ligands are highlighted in red. **(B)** Gene set enrichment analysis showing enrichment of the TGF-β signaling program in CD8⁺ T cells from liver metastases. The normalized enrichment score, nominal P value, and false discovery rate are shown. **(C)** UMAP projection and module score distributions showing enrichment of the TGF-β signaling program in CD8⁺ T cells from liver compared with non-liver metastases. **(D)** Combined TGF-â1 and TGF-â2 concentrations in regular activation medium and conditioned medium from DMS273 cells, HHL5 hepatocytes, or DMS273/HHL5 co-cultures (mean ± SEM, n = 3). ∗, P < 0.05; ∗∗, P < 0.01. **(E)** Immunoblot analysis of phosphorylated SMAD2, total SMAD2, TGF-β, and TGF-β receptor II in CD8⁺ T cells cultured in regular activation medium, the indicated conditioned media, or medium supplemented with 10 mmol/L lactic acid. α-Tubulin served as the loading control. **(F)** ChIP-qPCR analysis of H3K18 lactylation enrichment at the TGFB1 promoter in CD8⁺ T cells cultured with DMS273-conditioned medium, DMS273/HHL5 co-culture-conditioned medium, or 10 mmol/L lactic acid. Bars represent the mean of two biological replicates (n = 2), with individual biological replicates shown. **(G)** Time-course analysis of CD8⁺ T cell proliferation during culture with DMS273/HHL5 co-culture-conditioned medium in the presence or absence of a TGF-β receptor inhibitor. Data are presented as mean ± SEM. **Abbreviations:** AUPR, area under the precision-recall curve; FDR, false discovery rate; NES, normalized enrichment score; TGF-β, transforming growth factor beta; TGF-β RII, transforming growth factor beta receptor II; TGF-βi, transforming growth factor beta receptor inhibitor LY3200882

Lactate-associated scores also correlated positively with TGF-β-associated dysfunction signatures across patient samples, with the strongest relationship observed in liver-derived tumors (**Supplementary Fig. S5A**). This association suggested that lactate-related metabolic stress and TGF-β signaling are coordinated within the hepatic metastatic microenvironment.

The conditioned-media model was then used to examine the source and activity of TGF-β. DMS273/HHL5 co-culture medium contained higher concentrations of TGF-β1 and TGF-β2 than medium conditioned by either DMS273 or HHL5 cells alone **(Fig. 5D**), indicating that tumor–hepatocyte interactions generate a paracrine TGF-β signal.

Exposure of CD8⁺ T cells to DMS273/HHL5-conditioned medium increased TGF-β, TGFβRII, and SMAD2 phosphorylation, consistent with activation of canonical TGF-β signaling (**Fig. 5E**). Lactate supplementation produced a similar increase in these pathway components, suggesting that lactate can also engage the TGF-β program in recipient T cells.

Because lactate accumulation was associated with histone lactylation at exhaustion-related loci, H3K18la occupancy at the TGFB1 promoter was assessed by ChIP–qPCR. DMS273/HHL5-conditioned medium increased H3K18la enrichment at the TGFB1 promoter, exceeding that observed with lactate supplementation (**Fig. 5F**). This finding raises the possibility that lactate-associated histone lactylation contributes to the induction or maintenance of TGF-β signaling in CD8⁺ T cells.

The functional contribution of this pathway was tested using the TGF-β receptor inhibitor LY3200882. DMS273/HHL5-conditioned medium markedly reduced CD8⁺ T cell expansion, whereas TGF-β receptor inhibition restored proliferative capacity over time (**Fig. 5G**). Thus, paracrine TGF-β and lactate-associated activation of the TGF-β pathway converge to suppress CD8⁺ T cell proliferation in the SCLC hepatic metastatic niche.

### The lactate-TGF-β axis is associated with poor survival in SCLC liver metastases

The clinical relevance of these immune-metabolic programs was assessed using tumor transcriptional data from the phase III IMpower133 trial. Exhaustion, hypoxia, lactate, and TGF-β pathway scores were evaluated in Cox proportional hazards models adjusted for relevant clinical covariates, both in the overall cohort and after stratification by liver metastasis status.

Adverse associations with overall survival were most pronounced among patients with liver metastases. Exhaustion, hypoxia, lactate, and TGF-β signatures all showed higher hazard-ratio estimates in the liver-metastatic subgroup than in the overall cohort or in patients without liver metastases (**Fig. 6A**). Among the individual and composite measures, the combined lactate– TGF-β score showed the strongest adverse association with survival in patients with liver metastases (**Fig. 6B**).

**Fig. 6:**
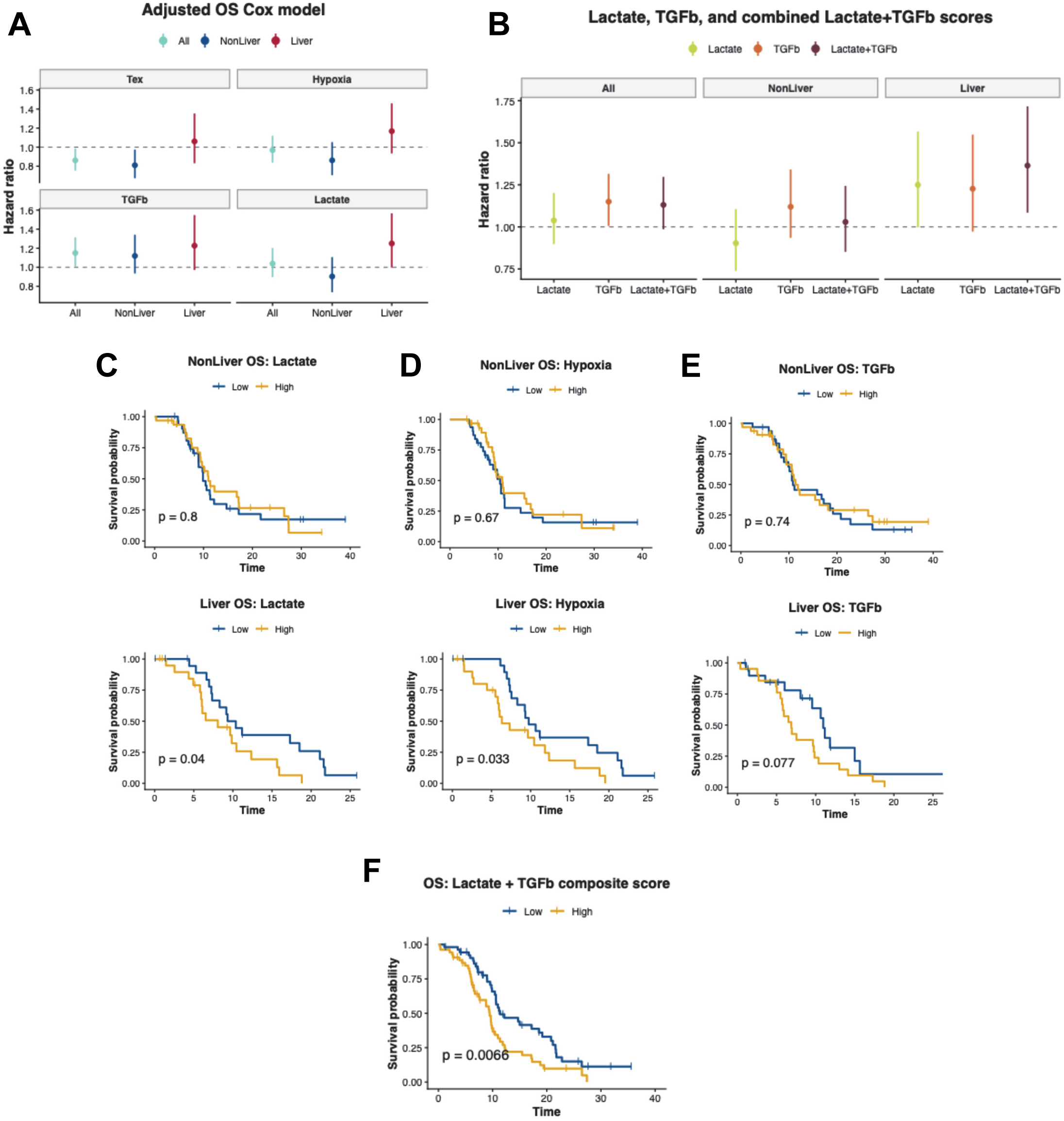
Coordinated lactate and TGF-β programs are associated with poor survival in SCLC. **(A)** Adjusted Cox proportional hazards models evaluating the associations of T cell exhaustion, hypoxia, TGF-β, and lactate signature scores with overall survival in the complete cohort and in patients with non-liver or liver metastases. Symbols indicate hazard ratios, and vertical lines indicate 95% confidence intervals. **(B)** Adjusted Cox proportional hazards models evaluating the associations of lactate, TGF-β, and combined lactate– TGF-β scores with overall survival in the complete cohort and in patients with non-liver or liver metastases. Symbols indicate hazard ratios, and vertical lines indicate 95% confidence intervals. In A and B, the dashed horizontal line denotes a hazard ratio of 1. **(C-E)** Kaplan-Meier analyses of overall survival according to low versus high lactate **(C)**, hypoxia **(D)**, and TGF-β **(E)** signature scores, shown separately for patients with non-liver metastases (top) and liver metastases (bottom). **(F)** Kaplan-Meier analysis of overall survival according to low versus high combined lactate-TGF-β composite scores. In C-F, P values were determined by log-rank test and are shown in the corresponding plots. **Abbreviations:** CI, confidence interval; HR, hazard ratio; OS, overall survival.

Kaplan–Meier analyses showed a similar pattern. Lactate, hypoxia, and TGF-β scores did not substantially stratify survival in patients without liver metastases (**Fig. 6C-E**). In contrast, high lactate and hypoxia scores were associated with significantly shorter survival in the liver-metastatic subgroup, while high TGF-β scores showed a similar adverse trend. A high combined lactate–TGF-β score was also associated with significantly shorter overall survival among patients with liver metastases (**Fig. 6F**). These clinical associations support the preferential relevance of coordinated lactate-related stress and TGF-β signaling in liver-metastatic SCLC.

## Discussion

Liver metastases represent a major clinical barrier to effective immunotherapy, yet the mechanisms of hepatic immune suppression remain incompletely understood. This study identifies a lactate–TGF-β axis linking SCLC–hepatocyte interactions to CD8⁺ T cell dysfunction. Across clinical trial cohorts, liver metastases defined a poor-prognosis subgroup with attenuated benefit from immune checkpoint blockade. Patient-derived single-cell and spatial analyses revealed exhausted CD8⁺ T cell states associated with hypoxia, lactate, and TGF-β signaling. Mechanistically, tumor–hepatocyte crosstalk generated convergent metabolic and cytokine signals that impaired T cell function, promoted histone lactylation at exhaustion-associated loci, and were partially reversed by TGF-β receptor inhibition. These findings extend prior work emphasizing reduced T cell abundance and macrophage-mediated T cell elimination in liver metastases (1–3,43) by identifying a complementary mechanism in which the hepatic niche imposes metabolic and cytokine stress on infiltrating CD8⁺ T cells. The association of combined lactate and TGF-β programs with inferior survival supports the clinical relevance of this immune-metabolic circuit.

Lactate emerged as a central component of this immune-metabolic phenotype. Tumor-derived lactic acid is known to suppress T- and NK-cell function (10,11,36). Our findings extend these observations to SCLC liver metastases by showing that interactions between tumor cells and hepatocytes generate more extracellular lactate than either cell type alone. Uptake of this lactate by recipient CD8⁺ T cells was supported by increased MCT1/SLC16A1 expression and stable-isotope tracing of co-culture-derived lactate into the T cells. Because histone lactylation links intracellular lactate availability to transcriptional regulation (41), its increase in recipient T cells provides a potential mechanism for sustained dysfunction. H3K18la was enriched at the PDCD1 and LAG3 promoters, and lactate supplementation reproduced this pattern, linking lactate exposure to an exhaustion-associated chromatin state. H3K18la enrichment at the TGFB1 promoter further raises the possibility that lactate also reinforces an immunosuppressive cytokine program in CD8⁺ T cells.

TGF-β emerged as a second component of this suppressive program. Unbiased ligand-activity analysis identified TGF-β as a candidate upstream regulator of the liver-enriched CD8⁺ T cell state, consistent with its established roles in hepatic immune tolerance, liver metastasis, and suppression of antitumor T cell immunity (5,12,13,42). The data point to two complementary sources of pathway activation: lactate exposure induced TGF-β signaling within CD8⁺ T cells, while SCLC–hepatocyte interactions increased extracellular TGF-β1 and TGF-β2. Both signals converged on canonical SMAD2 activation in recipient T cells. TGF-β receptor inhibition restored proliferation in the presence of SCLC–hepatocyte-conditioned medium, confirming a functional role for this pathway. These findings suggest that lactate-associated stress engages a T cell-intrinsic TGF-β program that is further amplified by paracrine TGF-β from the tumor– hepatocyte niche.

These findings have several translational implications. First, the combined lactate–TGF-β program may identify patients with liver-metastatic SCLC who are unlikely to benefit from immune checkpoint blockade alone. Second, restoration of T cell proliferation by TGF-β receptor inhibition provides a rationale for combining TGF-β pathway blockade with immunotherapy in this population. Strategies that reduce tumor lactate production or transport may provide an additional means of disrupting this circuit. Because systemic inhibition of lactate metabolism or TGF-β signaling can produce substantial toxicity, approaches that selectively target the tumor or hepatic metastatic niche may offer a more favorable therapeutic window. The combined signature could also provide a pharmacodynamic biomarker for evaluating whether such interventions effectively remodel the hepatic immune microenvironment.

Several limitations warrant consideration. Our conditioned-media system modeled soluble tumor–hepatocyte interactions but did not capture the full cellular and structural complexity of the hepatic niche, including macrophages, endothelial cells, stellate cells, and extracellular-matrix signals. Acute induction of PD-1 and LAG3 indicates an exhaustion-like state but does not by itself establish stable T cell exhaustion. Finally, our data associate histone lactylation with exhaustion-related transcriptional loci but do not establish that lactylation is required for their expression or for T cell dysfunction. Genetic or pharmacologic disruption of lactate transport and histone lactylation, followed by validation in immunocompetent models of SCLC liver metastasis, will be important next steps.

In conclusion, our study defines a hepatic immune-metabolic circuit in which SCLC–hepatocyte interactions generate lactate and TGF-β signals that converge on CD8⁺ T cells. Lactate accumulation accompanies histone lactylation at exhaustion- and TGFB1-associated genes, while paracrine and T cell-associated TGF-β signaling reinforces proliferative dysfunction. By connecting the clinical resistance phenotype of SCLC liver metastases to a targetable immune-metabolic mechanism, these findings provide a rationale for therapeutic strategies that disrupt the lactate–TGF-β axis to restore antitumor immunity.

## Data Availability

All data supporting the conclusions of this study are included in the main text, supplementary materials, and supplementary tables. Materials used in this study may be made available by the corresponding author upon reasonable request and in accordance with institutional policies and any applicable material transfer agreements.

## Authors’ Contributions

Conceptualization: A.K., A.T.

Methodology: A.K., Y.C., C.M., A.J., Y.Z., Y.H., C.T., B.S., T.A., A.K.S., A.T.

Investigation: A.K., Y.C., C.M., A.J., Y.Z., Y.H., C.T., B.S., T.A., A.K.S.

Visualization: A.K., Y.C., C.M., A.J., A.T. Funding acquisition: A.T.

Project administration: A.K., A.J., A.K.S., A.T.

Supervision: A.J., A.K.S., A.T. Writing-original draft: A.K., A.J., A.T.

Writing-review and editing: All authors.

## Supporting information

Supplementary Figures S1 through S5

## Acknowledgements

We thank the CCR Single Cell Analysis Facility for support with single-cell sequencing workflows and the NIH High-Performance Computing Biowulf cluster for providing computational resources and support. AI-assisted language tools were used to improve grammar, clarity, and readability.

