## Supplementary Figures S1 through S5 for "Tumor-hepatocyte crosstalk drives a hepatic lactate-TGF-β axis of CD8^+^ T cell exhaustion and immunotherapy resistance in small-cell lung cancer liver metastases"

### Supplementary Fig. S1

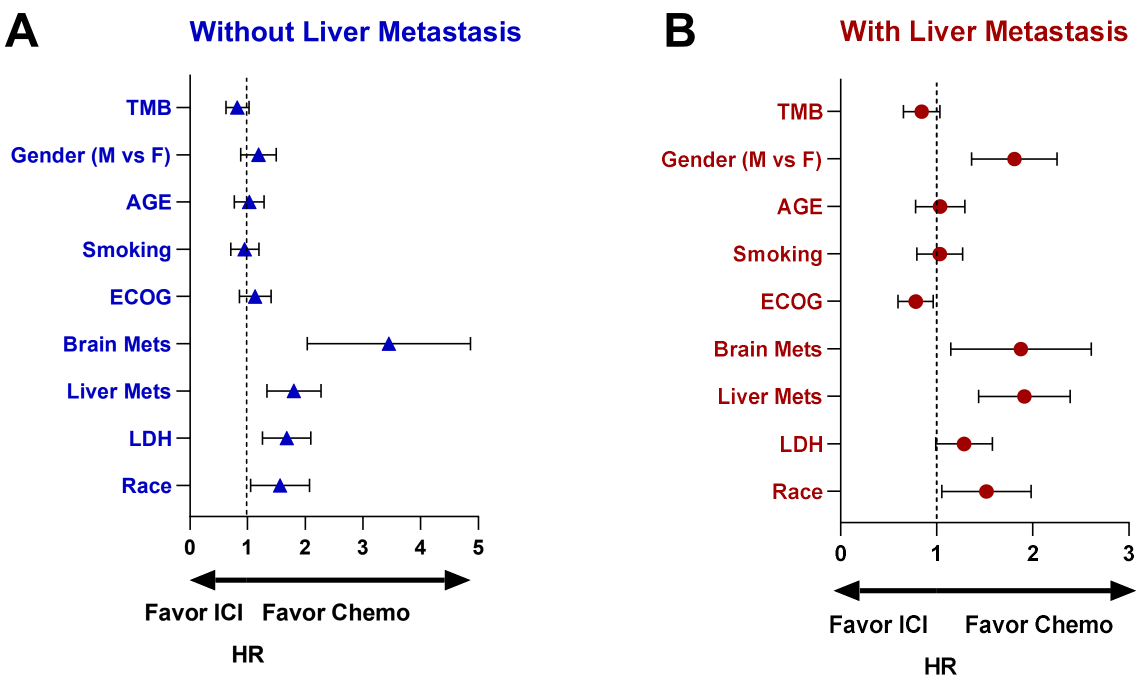

### Supplementary Fig. S2

**A**

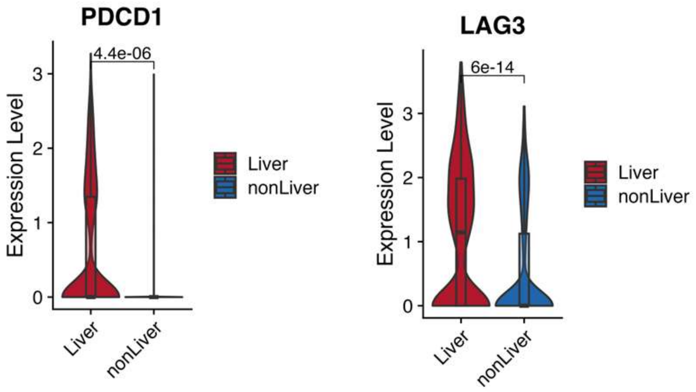

### Supplementary Fig. S3

A

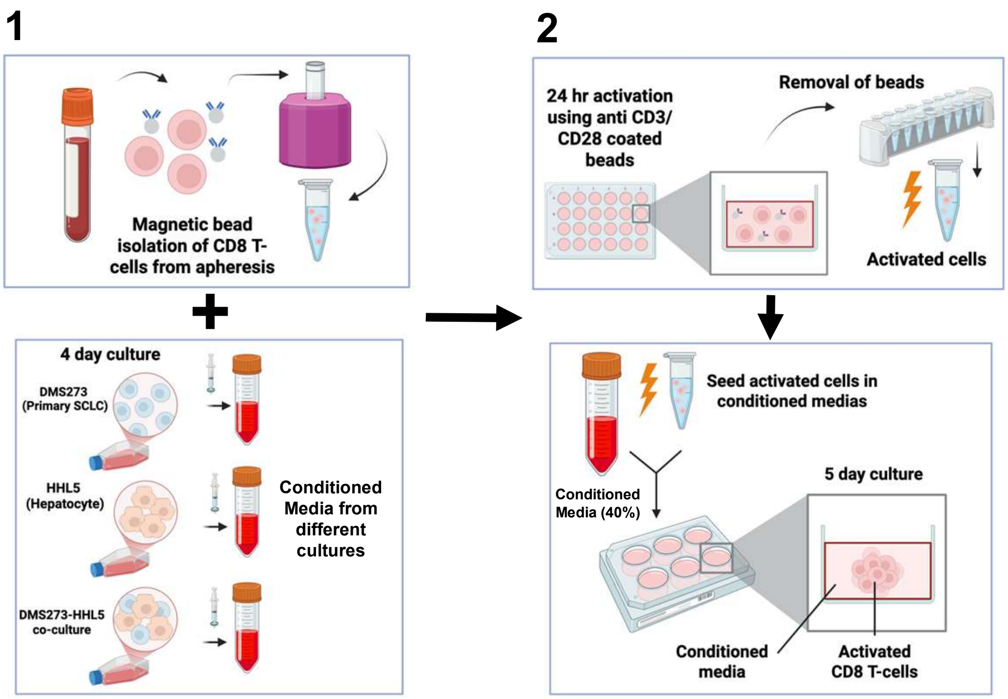


B

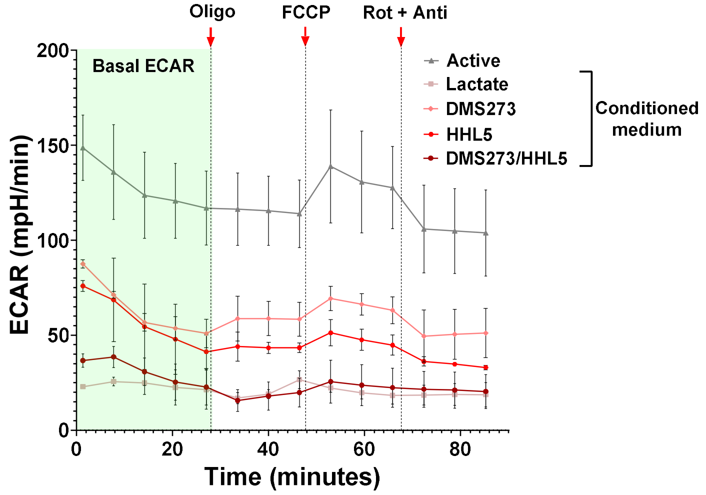

### Supplementary Fig. S4

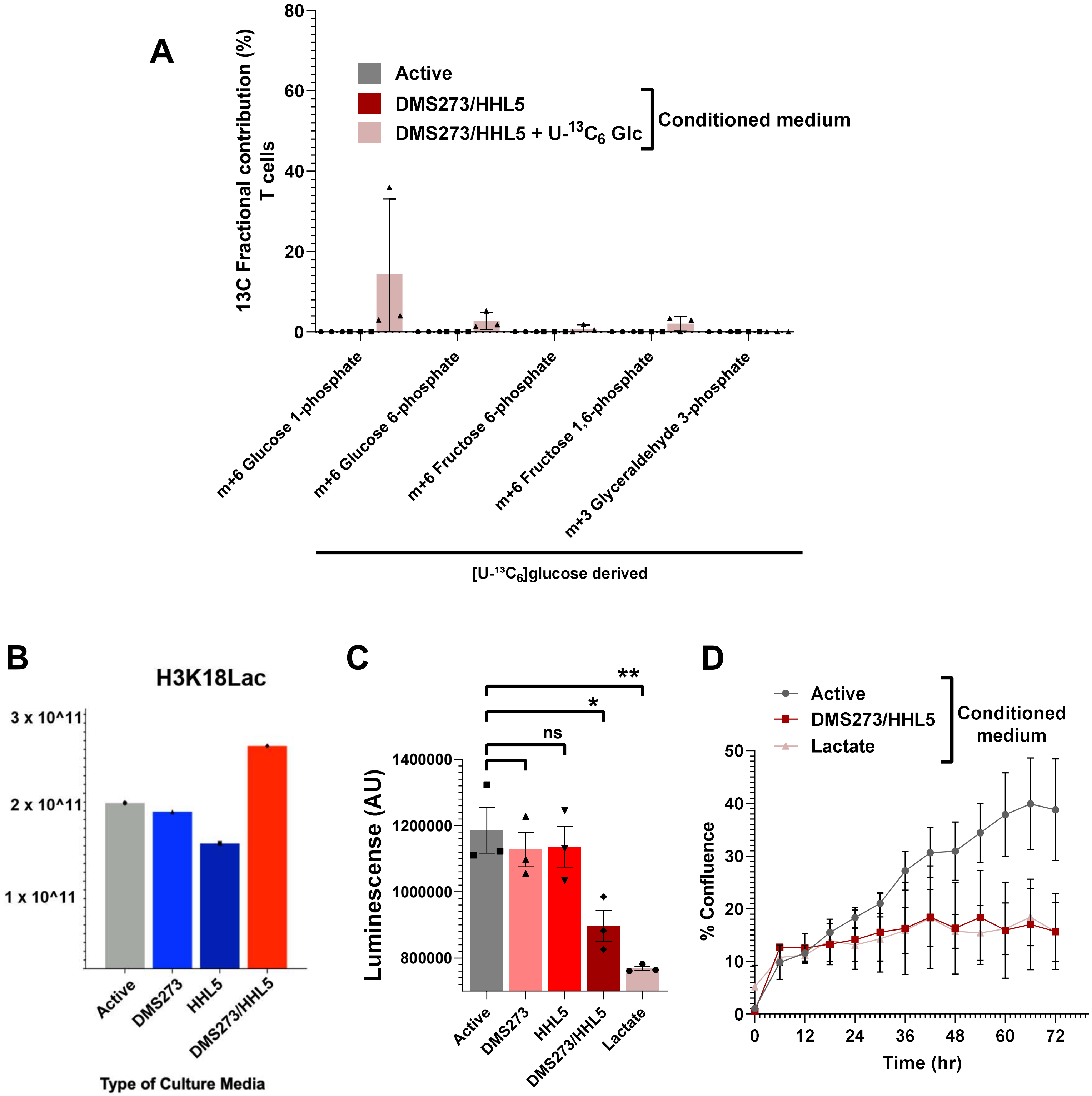

### Supplementary Fig. S5

A

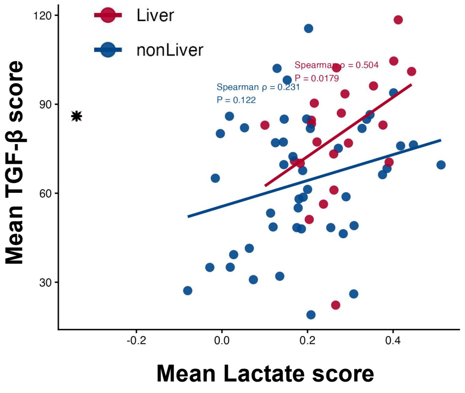
